# Flexible navigation with neuromodulated cognitive maps

**DOI:** 10.1101/2025.09.04.674155

**Authors:** Krubeal Danieli, Mikkel Elle Lepperød

## Abstract

Animals develop specialized cognitive maps during navigation, constructing environmental representations that facilitate efficient exploration and goal-directed planning. The hippocampal CA1 region is implicated as the primary neural substrate for cognitive mapping, housing spatially tuned cells that adapt based on behavioral patterns and internal states. Computational approaches to modeling these biological systems have employed various methodologies. Although labeled graphs with local spatial information and deep neural networks have provided computational frameworks for spatial navigation, significant limitations persist in modeling one-shot adaptive mapping. We introduce a biologically inspired place cell architecture that develops cognitive maps during exploration of novel environments. Our model implements a simulated agent for reward-driven navigation that forms spatial representations online. The architecture incorporates behaviorally relevant information through neuromodulatory signals that respond to environmental boundaries and reward locations. Learning combines rapid Hebbian plasticity, lateral competition, and targeted modulation of place cells. Analysis of the capabilities of the model on a variety of environments demonstrates our approach’s efficiency, achieving in one shot what traditional RL models require thousands of epochs to learn.

The simulation results show that the agent successfully explores and navigates to the target locations in various environments, showing adaptability when the reward positions change. Analysis of neuromodulated place cells reveals dynamic changes in neuronal density and tuning field size after behaviorally significant events. These findings align with experimental observations of reward effects on hippocampal spatial cells while providing computational support for the efficacy of biologically inspired approaches to cognitive mapping.

## 1 Introduction

Survival in complex environments requires efficient navigational strategies. From desert ants to humans, successful wayfinding—navigating toward goals that are not directly visible depends on emergent internal spatial representations, known as cognitive maps [1, 2]. Understanding how these maps are constructed from ongoing experiences and how they can be exploited for flexible goal-directed navigation remains an active area of research in both neuroscience and reinforcement learning (RL).

The hippocampus (HP) and the entorhinal cortex (EC) serve as the main neural substrates for spatial representation in the brain. They contain specialized neurons that encode spatial and contextual information, including grid, border, speed, and place cells [3, 4, 5]. Place cells in the CA1 hippocampal region have attracted particular interest due to the convergence of inputs from entorhinal grid cells, the CA3 hippocampal region, and the lateral EC [6, 7, 8, 9]. This strategic integration of diverse spatial and contextual signals suggests that CA1 place cells may play a critical role in the formation and maintenance of cognitive maps [10].

Traditional theories of cognitive maps suggest that spatial representations can emerge from multiple navigational strategies. One of these is path integration, which involves tracking one’s position by integrating past movements—using action or velocity vectors—to estimate the trajectory and return to the point of origin.

An important strength of path integration lies in its independence from external cues, instead relying on idiothetic signals, internal motion signals, considered biologically plausible and believed to involve entorhinal grid cells [11, 12]. A concrete implementation of this strategy is route learning, in which sequences of actions and positions are encoded along traveled paths. However, an approach that can scale poorly [13, 14, 15]. In contrast, survey maps are based on Euclidean geometry, offering more flexible navigation capabilities [14, 16] but at times with not biologically plausible geometric assumptions [17, 18, 19, 20]. A compromise is offered by labeled graph representations, which encode landmarks and transitions in a topological network. These support vector-like operations, planning, and prediction while avoiding strict geometric constraints [21, 22, 23]. On the computational side, various models have attempted to formalize and replicate these navigational strategies. Early frameworks suggested that the hippocampus encodes both spatial location and direction [24], while graphbased models captured structural aspects of spatial organization, although often at the cost of scalability [15]. Several studies focused more directly on path integration, typically using recurrent neural networks combined with linear readouts. These systems, when trained on navigation tasks, have spontaneously developed spatially-tuned activity patterns reminiscent of grid, place, and border cells [25, 26, 27].

Another proposal is the Tolman-Eichenbaum machine (TEM), which is generalized in spatial and relational tasks while mimicking biological neural activity patterns [28, 29, 30]. Learning occurs through gradient descent over many episodes or epochs, far away from adaptive and context-dependent learning seen in biological systems. Nevertheless, as most computational approaches these models possess simplified neuronal dynamics, often reduced to artificial neurons defined as a weighted sum and a fixed non-linear activation function.

An alternative is the successor representation framework, built around the learning of a predictive map of the explored state space, which can be used for active navigation [31, 32]. It has been associated with the hippocampus for the generation of a cognitive map, using the possibility of incorporating reward information and formulation in terms of biologically plausible mechanisms [33, 34, 35, 36].

Another relevant element in brain dynamics are neuromodulators, endogenous molecules with a range of specialized actions that affect physiology, cognition, and behavior [37]. Their functions include filtering meaningful internal and external signals, influencing neuronal dynamics, and learning by altering synaptic states and parameters [38, 39, 40]. An important neuromodulator is dopamine, long associated with reward information, encoding of prediction errors [41, 40] and novelty detection [39], mechanisms closely related to reinforcement learning principles [42, 43]. In addition, projections from the Ventral Tegmental Area (VTA) and Locus Coeruleus (LC) have been shown to target the hippocampal circuits [44, 45, 46], to be involved in memory formation [47], and affect the adjustment of CA1 places cells [48, 49, 50, 51, 49], particularly through LEC inputs [52, 53].

Others incorporate reward-driven Hebbian plasticity modulated by neuromodulators [54]. Nevertheless, these architectures mostly fail to unify these ingredients into a biologically grounded system that at the same time learns a map of the environment online without relying on an external coordinate system, and flexibly performs goal-directed navigation.

In this work, we present a biologically inspired model of cognitive map formation that integrates grid cells, place cell representations, neuromodulatory signals, and graph-based spatial computations. We first aim is to demonstrate that an agent endowed with such bioinspired model architecture is capable of building content-rich topological maps of the surroundings on the fly, and leveraging it for efficient goal-directed navigation. The testing protocol was a common behavioral and reinforcement learning task: exploration of a novel environment and the localization and collection of a reward. Further validation was performed in multiple experiments in which some aspects of the protocol were varied.

For building the cognitive map, a strategy similar to path integration has been used in that the agent develops a place cells representation as it navigates in unexplored spaces. However, spatial tuning is achieved through fast synaptic plasticity and competition, and not through numerous backpropagation cycles. The use of idiothetic information only and not other sensory modalities, such as visual cues, is motivated by reducing architectural complexity and proving the validity of the approach with minimal sources of information.

Concerning neuromodulation, it has been operationalized as a collection of variables acting as low-pass filters of sensory events [55, 41, 35], here being only rewards and collisions with boundaries. The action of neuromodulators is to form adaptive connections with place cells through local Hebbian plasticity, effectively determining a scalar field over the spatial map [56] that can be used for planning [57, [58, [59, [60, [29]. Moreover, they were also given the opportunity to directly modulate some neuronal properties of place cells, such as the gain of neural activation and the place field.

In fact, another objective of ours was to evaluate how active neuromodulation of neuronal dynamics can impact task performance. To this end, multiple comparative ablation experiments were performed to highlight the relevance of each modulatory mechanism.

Finally, we demonstrate how the model dynamically adapts to environmental changes and how some neuromodulatory actions can positively influence the formation and flexibility of a cognitive map [61, 62].

The remainder of the paper is organized as follows: Section 2 details the model and experimental setup; Section 3 presents results; Section 4 discusses broader implications and future directions.

## 2 Methods

We propose a model of cognitive map formation driven by an agent’s experience within a closed environment.

The architecture operates with minimal external inputs, limited to binary reward and collision signals, as illustrated in Figure 1**a**. Instead of relying on exteroceptive cues, spatial representations emerge from idiothetic information, that is, the agent’s internal perception of self-motion [63], consistent with the integration frameworks of the prior path. In practice, we use the agent’s ground truth velocity vector, that is, its actual displacement within the environment, as the primary navigational signal, reflecting the integration of inertial and proprioceptive signals observed in biological systems [64, 11]. Since no visual information is used, the agent effectively navigates in the dark.

**Figure 1.**
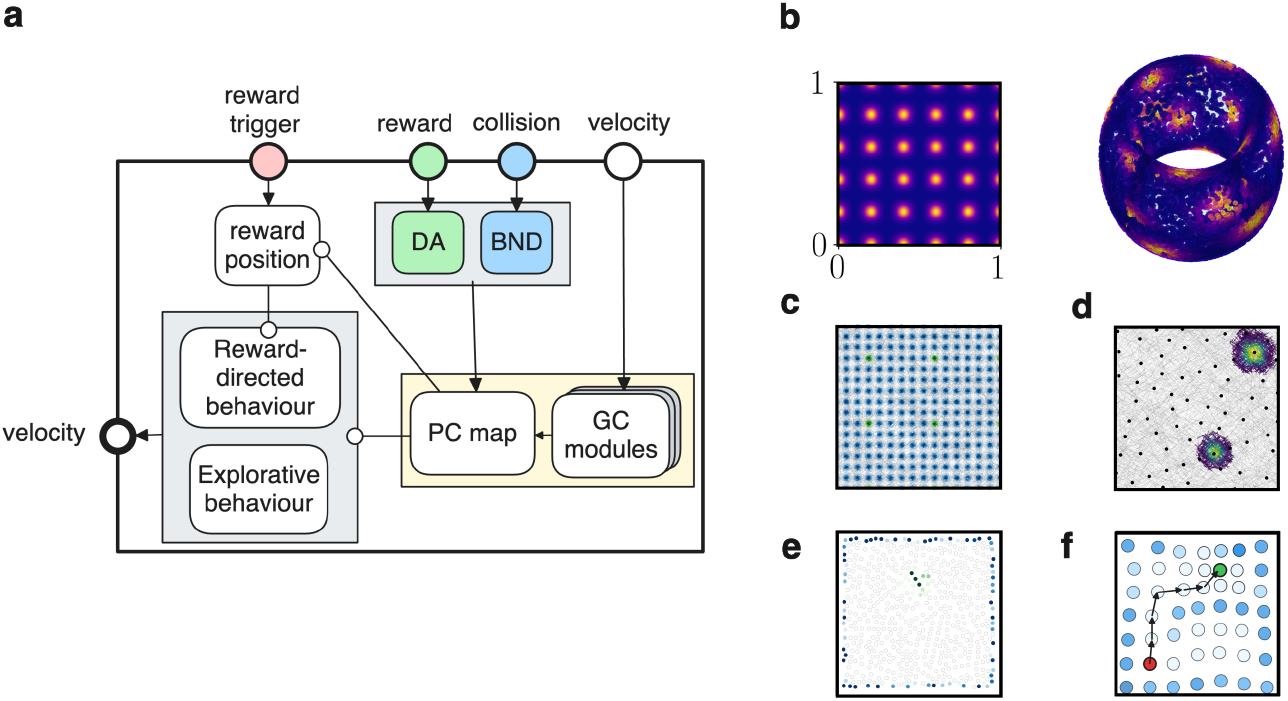
Model layout and spatial representations. **a**: the full architecture of the model, consisting of three main sensory inputs, targeting the two modulators and the cognitive map module, and the executive components, represented by a policy module, two behavioral programs, and a reward receiver. **b**: a module of grid cells defined in a bounded square space of length 1, and an activity representation of their receptive field over a torus. **c**: the neural activity of a grid cell module from a random trajectory; in blue the repeating activity of all cells, while in green the activity of only one, highlighting the periodicity in space. **d**: the distribution in space of the place cells centers, together with the activity of two cells showing the size of their place field. **e**: neuromodulation activity over the place cells map, with in blue the cells tagged by the collision modulation, and in green the ones targeted by reward modulation. **f**: The place cells layer can be regarded as a graph with values assigned to each node according to the modulation strength; a path-finding algorithm can then be used to connect any two nodes, taking into account the node values.

### Place Cell Formation

The primary spatial representation is formed by a set of simplified grid cell modules, each encoding a periodic tiling of 2D space, which directly maps to a toroidal manifold **T**^2^ (Fig. 1**b**). Leaving traditional grid cell modeling approaches [65, 66], we generate population activity directly by Gaussian tuning on a torus, continuously updated using the agent velocity vector, an approach used in previous work [8].

The grid cell population vector **u**^GC^ is forwarded to a place cell network with initial zero synaptic weights. When no place cell is sufficiently active for a given input, a silent unit is randomly selected and imprinted with the current grid activity pattern. To enforce sparsity in the population vector of place cells and clear spatial tuning specificity, a lateral inhibition mechanism is implemented by comparing the cosine similarity between the new weight vector 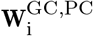 and the existing non-zero weights 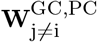 against a threshold 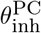, if it is above then the new place cell is discarded.

The activation of each place cell is calculated through a bounded cosine similarity function, determining its corresponding place field (Fig. 1**d**). Further implementation details, including lateral inhibition and recurrent connectivity, are provided in the appendix.

### Neuromodulation

Neuromodulators deliver event signals: rewards, denoted DA (for dopamine), and boundary collisions, denoted BND. In particular, the latter is an internal variable that correlates and tracks the sensory experience of mechanical collisions. In the brain, this functionality might be partially supported by serotonin (5-HT), which has been proposed to influence and affect sensory processing [67, 68], encode unexpected uncertainty [69], modulate neural excitability and synaptic plasticity [70, 42, 71], also in CA1 neurons in the hippocampus [72, 73].

They are driven by binary inputs and defined through a leaky variable with exponential decay.

Each modulator k updates its connection weights with the place cells through a Hebbian rule based on place cell activity **u**^PC^ and the error:

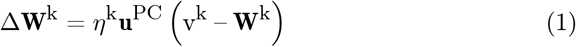

The term in brackets can be considered an error, implementing a simple form of predictive coding, and is inspired by temporal-difference learning [43], aligning with evidence that neuromodulatory systems signal prediction errors and update beliefs [74, 29, 75]. This plasticity mechanism provides resilience to environmental changes (e.g., moving rewards and new walls).

The weight vectors are restricted to remain non-negative. Reward modulation tags cells near the rewarded locations, whereas boundary modulation builds a representation of the environmental edges. The result is a set of scalar fields over the place cells, and it forms the core of our model of a cognitive map (Fig. 1**e**). See the appendix for full learning rules and parameter settings.

### Modulation of Place Fields

In addition, we tested whether neuromodulators could directly alter spatial tuning. Place fields were dynamically shifted and resized based on recent salience signals.

Following a salient event (reward or collision), the place field centers were displaced in the grid cell space with the magnitude scaled by the neuromodulator v^k^ and proximity to the event:

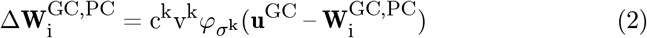

Here, *φ* is a Gaussian function with width *σ*^k^ and c^k^ a scaling factor. This rule is inspired by BTSP plasticity [50, 61], which shifts CA1 place fields following salient experiences. This action was applied only to recently active cells, that is, with an activity trace greater than a threshold *θ*^k^. In addition, lateral inhibition prevents field overlap during re-mapping.

In addition to dislocation, the field size was modulated by scaling the gain of recently active neurons. This mechanism allows neuromodulators to transiently enhance or suppress spatial sensitivity for specific cells. This modulation rule involves the gain *β*_i_ of each cell being adjusted proportionally to its activity trace m_i_, a reference gain constant 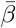, and a modulatory scaling variable:

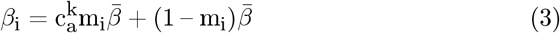

where 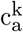 is a scaling gain parameter, for which a value of 1 means that there is no modulation.

### Policy and Behavior

To evaluate the utility of the model in navigation, we implemented a simple policy that toggled between exploration and reward-seeking behavior, depending on an external reward trigger and the internal map.

Planning was defined as the action protocol of selecting a goal position; the calculation of a trajectory as a sequence of place cells starting with the one corresponding to the current position and ending with the nearest to the goal position; and the execution of actions to visit the entire sequence. For the trajectory, a graph-based pathfinding algorithm was used, in which place cells served as nodes, while synaptic connections acted as edges. In addition, a cost function was introduced in the graph nodes, which assigns lower values to BND-modulated cells. This was designed to discourage proximity to the walls of the environment, as shown in plot 1**f**.

Exploration consisted of two possible strategies: a random walk, for purely stochastic movements, and periodic goal-directed navigation towards a random but visited location, aimed at preventing stagnation.

In contrast, exploitation, defined as reward-directed navigation, involved identifying the location of the reward within the cognitive map. This location corresponded to the average position of DA-modulated place cells, reflecting mechanisms such as hippocampal replay and value-based navigation [76, 77, 78].

### Naturalistic task

The model was evaluated on a biologically inspired navigation benchmark involving exploration and goal-seeking behavior in closed environments. Performance was measured as the total number of rewards collected in multiple trials.

Optimization of the model hyperparameters was carried out using the evolutionary Covariance-Matrix Adaptation Strategy (CMA-ES) [79]. The choice of hyperparameters to evolve took into account the type of dynamics for which it was difficult to select predefined values; the number of evolved variables was 10. The use of such method derived from the impracticality of other optimization algorithms, also given the non-differentiability of several dynamic. Viable alternatives were based on reinforcement learning, Bayesian, and grid search, but evolution was more appealing for the exploration and visualization of parameter space exploration. For evaluating the individuals, a multi-objective fitness function was used: maximization of the reward count, and minimization of the collision count from the time the reward position was discovered.

## 3 Results

### Performance in wayfinding

Our primary aim was to evaluate the formation of the cognitive map through neuromodulation in terms of the performance of the goal navigation in different environments. The best model resulting from evolution reached solid navigation and adaptation skills. The agent was able to visit a significant portion of the environment during exploration and use neuromodulation to produce useful spatial representations.

The left panel of Fig. 2**a**-**b** displays place cells associated with collisions and reward events, signaling boundaries (blue) and reward (green) locations. The overlap of these two representations and the non-modulated place cells (in gray) is what we refer to as a cognitive map, since these are the main sources of spatial and contextual information used during planned navigation, whose path is depicted as a gray line. The right panel instead portrays the actual environment with walls (black), reward location (green), and multiple trajectories (red). During exploration, the main areas were visited until the reward position was located and the goal-directed navigation dominated, as highlighted by the density of the path lines. Considering the position of the walls and corners, the layout of this environment does not always make the target locations visible, as it is a non-convex area and therefore can be classified as wayfinding [80]. The challenge of not being able to use straight lines is overcome by the graph approach using local data and the consideration of boundary place cells, allowing the agent to plan accordingly. In addition, considering the Gaussian receptive fields and the approximately homogenous distribution of place field centers supported by lateral inhibition, the calculated path accounting for node-length also implicitly minimizes effective path-length, although not necessarily exactly. Figure 2**i** visualizes part of the path-finding process.

**Figure 2.**
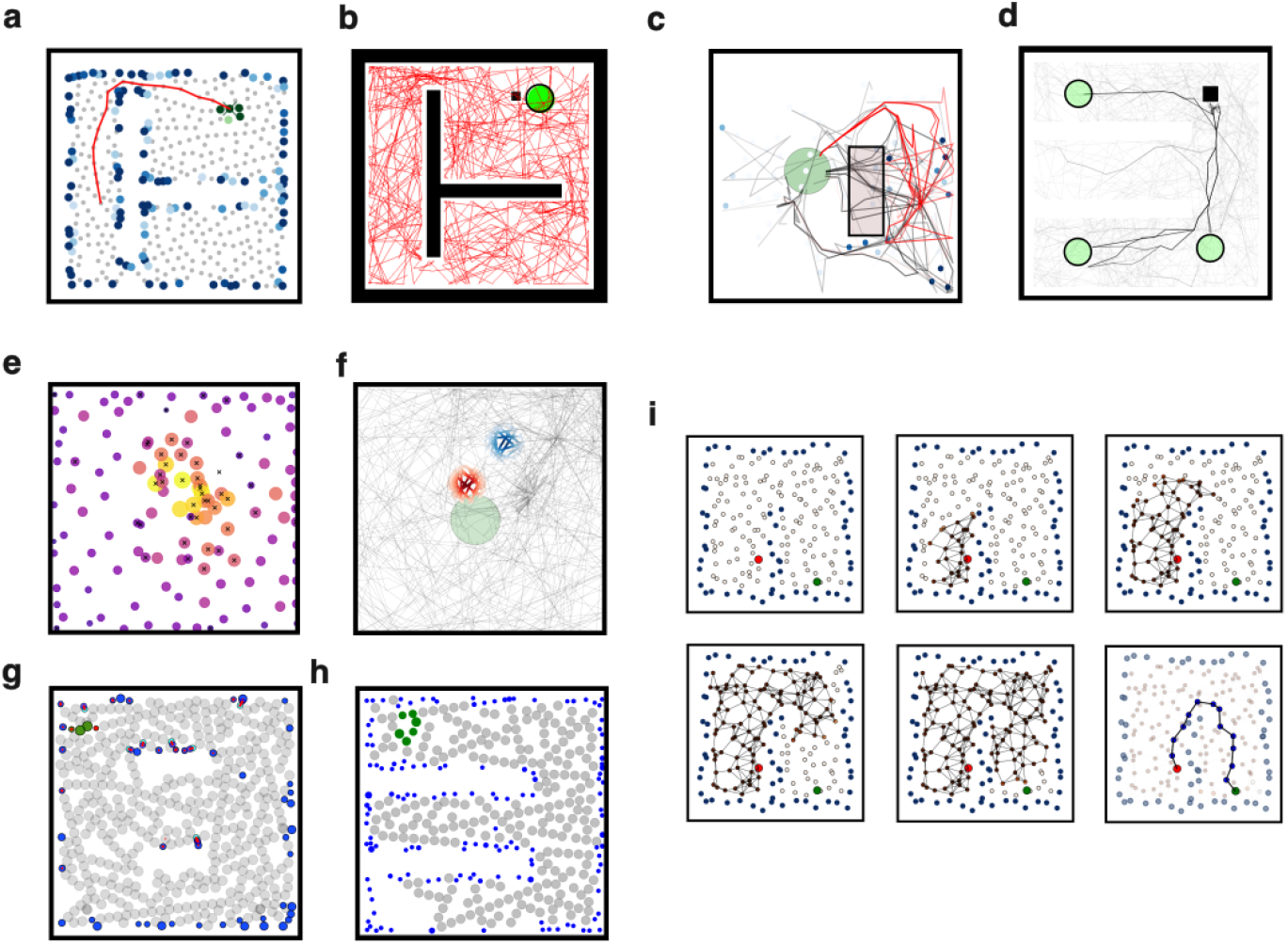
Cognitive maps and performance results. **a**: a cognitive map over a space, together with the plan (red line) to reach a target location from a starting position. **b**: the same environment but with the reward (green circle), trajectory (red line), agent position (black square). - **c**: plot of trajectories before (black) and after (red) the insertion of a wall (rectangle) between the starting and goal positions, the wall can also be spotted from the boundary cells in blue. - **d**: trajectories for multiple trials with the agent starting at the same position (black square) but with the reward location (green circles) periodically moving. - **e**: place cells centers with circle size and color proportional to their node degree; cells with a black cross changed their place field location over time. - **f**: place fields of the same cell before and after several relocation of its center following reward events. - **g**: place cells centers with size inversely proportional to their gain value; in blue boudary cells with the highest average gain, in green reward cells with second smallest gain, the others in grey; red lines represent the axis of relocation of the place fields. - **h**: similar plot for another environment, without relocation axes, - **i**: visualization of part of the path-finding algorithm, propagation of an activity wave through the place cells network from top-left to bottom-center, and the calculated path visualization in the bottom-right.

In general, this result confirms the ability of the model to focus on navigation and obstacle avoidance. However, it is worth noting that not all simulations resulted in a reward being found in the first place, due to the randomness of the exploratory process; this effect was more pronounced in environments with more walls and narrow passages.

### Detour task

The planning ability and plastic nature of the cognitive map should provide resilience against unexpected changes in the environmental layout. In order to verify this, we implemented a detour experiment. Initially, the agent was familiarized with a square environment with the reward in the middle and always starting from the same position. Then, a wall was placed between the starting position and the reward, therefore forcing a new trajectory to reach it. As expected, the agent was able to form a representation of the new obstacle and calculate new paths around it, succeeding in the task. In Fig. 2**c** they are shown the trajectories before and after the wall placement, and they manifest the ability of detour in the new layout.

### Adaptive goal representation through sensory error

Then, we tested the adaptability to environmental changes. In this scenario, the reward object was moved after being fetched a fixed number of times. Here, the difficulty was to unlearn previous locations and discover new ones, in a protocol similar to [54]. In Fig. 2**d** is reported the set of trajectories over many trials with the reward displaced in three possible locations. The agent was capable of planning behavior, as earlier, but also exploring and finding new rewards, as shown by the density of lines. Whenever a goal path resulted in a failed prediction, the DA-based sensory error weakened the association between the place cells and the reward signal, leading to an extinction of its representation at that location. This result validates the resilience of the model to changing sensory expectations, in this case the reward position.

### Modulation of place field size

The construction of the model is such that the experience of environmental events can impact the neuronal properties of the generated place cells. In particular, collision and reward events have the effect of affecting the neural activation gain *β* of BND and DA-modulated cells through a hyperparameter.

The hyperparameter values 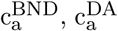, that yielded the best results were larger than 1, 4.6, 4.4, respectively, causing a decrease in field size. In Fig. 2**g**-**h** are showed the cognitive maps with relative place field sizes for two environments are shown, showcasing the differences between boundary, reward and nonmodulated cells. In addition, Fig. 2**h** indicates the place cells that underwent re-location of their fields with a red line representing the displacement vector.

The distribution of this re-location vector is biased towards the rewarding area, a similar observation to Fig. 2**e** in which the cells involved in the modulation of field positions are marked with a black cross. In Fig. 3**d** is showed the evolution of the gain magnitude for a sample of boundary cells and reward cells over their corresponding modulatory events. Notable is the possible decrease in value, a case that occurs when a cell is active but the modulation is absent, and thus the gain is pushed down towards the baseline 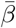. This feature can be considered another adaptation property.

**Figure 3.**
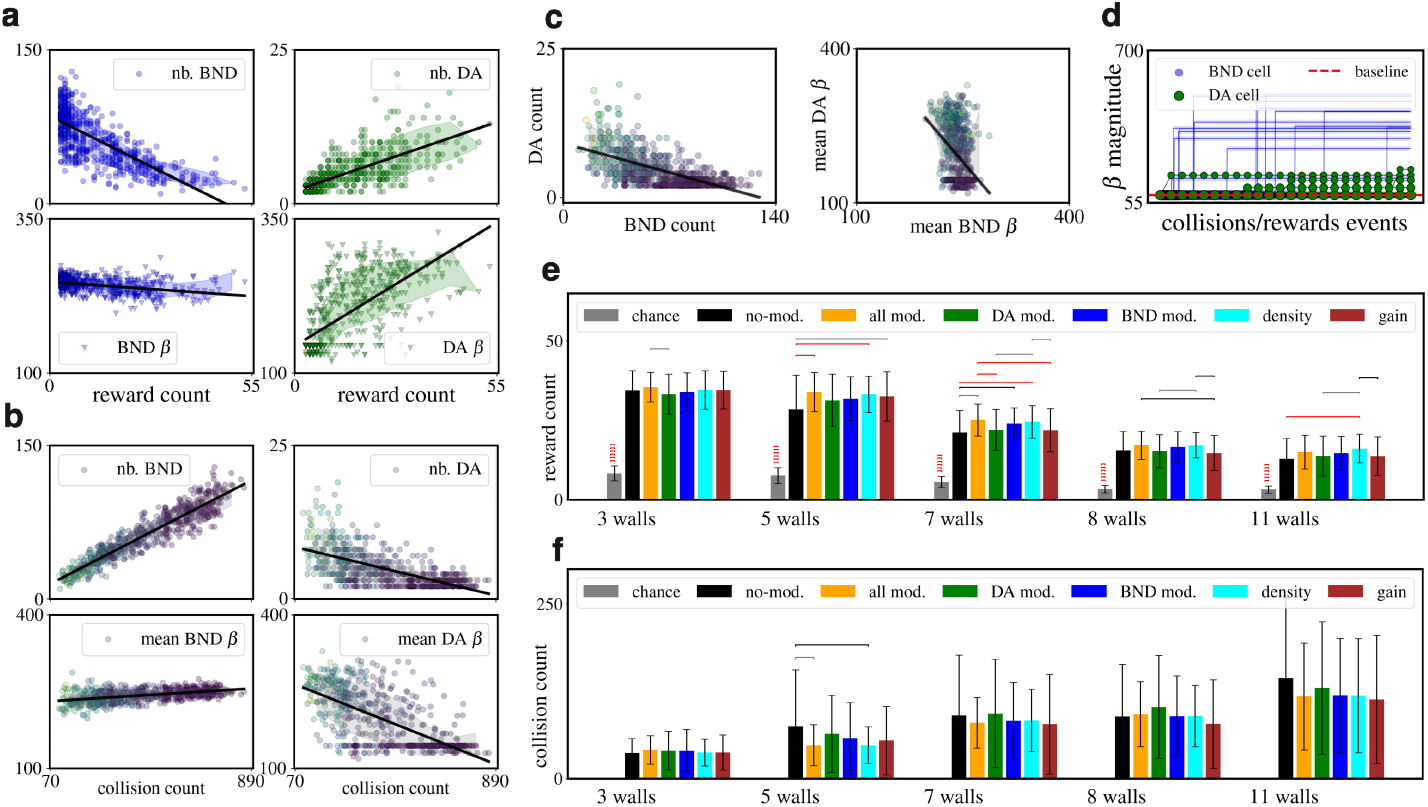
Cognitive maps and performance results. **a**: effect of reward count on reward and boundary modulated cells (green and blue respectively), both in total count (top row) and average gain modulation magnitude (bottom row); simulation of 160 independent runs. - **b**: similar plot but with respect to collision count and with a color map of the reward count - **c**: relation between count of reward and boundary modulated cells, and between gain modulation magnitude; color map of the reward count - **d**: gain magnitude of boundary and reward cells over sequences of collision and reward events respectively - **e**: reward count performance comparison for the same base model but with different levels of ablation: chance level with only random walks, completely without modulatory actions, all modulation enabled, only DA-related modulation, only BND-related modulation, only place cells density modulation, only activation gain modulation. Pair-wise t-test over 128 iterations and Bonferroni correction - **f**: collision count from the time a reward representation is formed (first non-zero DA weights), same ablation variants.

### Effect of modulation on performance

Lastly, we investigated the effect of modulating the density of place cells and the size of the field. The goal position was fixed, but the agent was randomly relocated after fetching; performance was defined as the total number of reward counts within a time window.

Our working hypothesis is that these experience-driven neuronal changes would improve the quality of the cognitive map and be reflected in navigational abilities. The assessment of this claim was conducted by comparing variants of the model, obtained by progressively ablating the reward (DA) and collision (BND) modulatory actions, as well as the density and gain mechanisms separately. All models were run in five different environments that differed by the number of internal walls, from 2 to 11, for a total of 128 independent simulation repetitions performed for each case.

The statistical results are shown in Fig. 3**e**. All models performed above chance, but the main finding is the confirmation of the importance of neuronal modulation, as revealed by the statistical difference of most modulation-powered models with respect to the one with all modulation disabled. In Fig. 3**f** is reported, for the same runs, the number of collisions from the time the reward is learned. Despite the large variance among the runs with the same simulation settings, the higher count is noticeable for the model without modulation. This finding highlights the role of boundary modulation in improving reward-directed behavior, possibly by changing the graph distribution around edges and corners to lower the total path length.

An additional finding was that density modulation is the main action behind behavioral improvements and alone does not perform worse than the model with all actions together.

Figure 2**d** showcases the distribution of place cells with the circle size and color representing the degree of the node, which is discretely aligned with the reward position and density. Furthermore, in Fig. 2**e** the place field of one cell is shown, before and after several reward occurrences and consequent center relocation.

Taken together, these findings support the hypothesis of the practical utility of direct modulation of place field structure for active navigation, even in these simple settings.

### Convergence of evolved hyperparameters

The use of an evolutionary algorithm for selecting the model dynamics had the convenience of providing a distribution of hyper-parameters over a population. In Figure 4, each point represents the value of one individual plotted with respect to its average reward count. A marked convergence is visible for most variables, with some displaying a quasi-bimodal distribution. The hyperparameters directly involved in neural activation, such as the parameters of the activation function *β, α*, and the trace time constant *τ* ^PC^, display an elevated degree of clustering. In addition, their specific values are such that the model is brought together to develop a dense place cell network with weak lateral inhibition given the high threshold 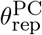 and low field overlap, due to the steep gain *β*.

**Figure 4.**
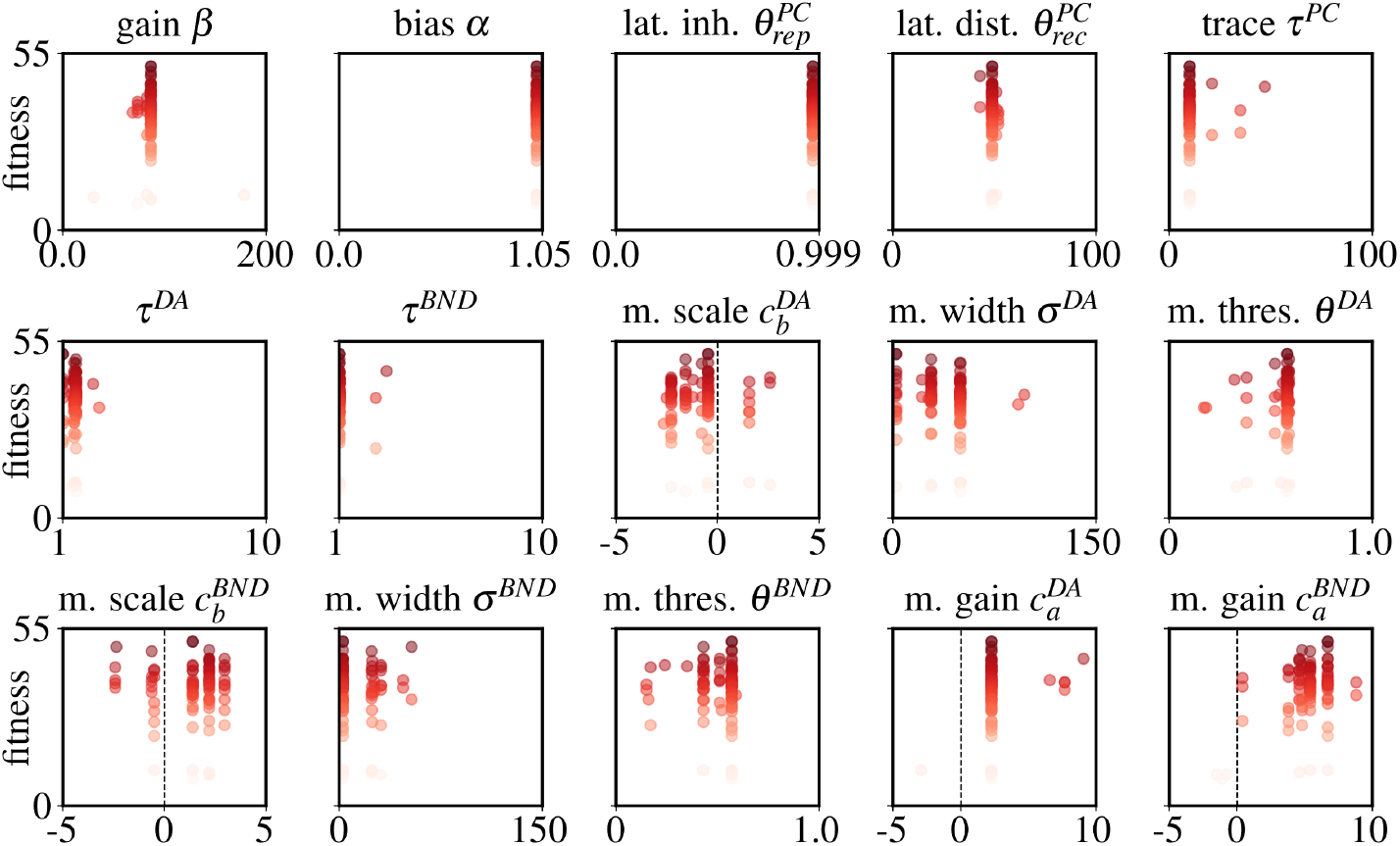
Distribution of evolved hyper-parameters. Results relative to the last generation, from a run with population size of 96 individuals. The hyper-parameters are: neural activation gain β, neural activation bias α, lateral inhibition threshold 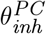, lateral distance threshold 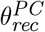, activity trace time con-stant τ ^PC^, reward DA modulation time constant τ ^DA^, boundary BND modulation time constant τ ^BND^, reward DA modulation scale 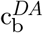, reward DA modulation spread 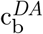, reward DA modulation threshold 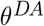, boundary BND modulation scale 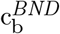, boundary BND modulation spread 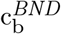, boundary BND modula-tion threshold θ ^BND^, reward DA gain modulation 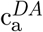, and boundary BND gain modulation 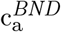.

In terms of modulation, there is a strong tendency to increase the magnitude of the gain for both reward 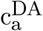 and collision 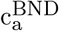 events, with the effect of further reducing the size of the active place fields. This result is in the direction of experimental observation showing a shrinkage of place fields near objects and walls [81, 82].

Instead, in modulating the centers of the place cells, there is more diversity in the action adopted. In fact, it can be both pulling towards and pushing away from the current agent position, with the latter being more frequent in regard to collision modulation. These findings align, at least partially, with previous work on remapping of place fields [83] and reward-induced changes in the preferred location for the position of hippocampal cells [84, 85, 86, 50].

In summary, over the course of generations, evolution converged to specific neural dynamics that are reminiscent of patterns observed in biological neuronal systems.

## 4 Discussion

Exploration and planning in novel and past environments are essential behaviors of animals that directly affect their success in spatial understanding and achieving goals.

An important element behind these abilities is the formation of a map of their surroundings, enriched with information gained from new experiences, known as a cognitive map. In this work, we presented a rate network model inspired by the CA1 hippocampal region [10], which differs from previous approaches as it operates online, uses neuromodulation-based plasticity and does not rely on external coordinates.

We used simplified grid cells together with synaptic plasticity as a mechanism to develop information-rich representations based on place cells updated through experience, grouped with common perspectives on cognitive maps [87]. In the spirit of minimizing geometric assumptions in the neural space, we treated the generated place network as a topological graph learned through velocity inputs, reminiscent of path integration [16, 88, 12], and with sensory information added locally through the action of neuromodulators. This idea aligned with the concept of a *labeled graph* [89, 18], however, it is also true that no metric violations were possible in these settings.

The tasks we applied the agent to consisted of an exploratory and exploitative phase, in which the system was tasked to plan and reach reward positions. For simplicity, the first stage relied on a random walk process, as it was outside the scope of this work. This choice had the side effect that the reward was not always discovered, leading to the formation of incomplete maps, and thus impairing performance. However, this issue had limited consequences.

The simulation results validated the model, showing the expected emergence of cognitive maps and their encoding of information collected during the experience. Previous work has used path integration with deep neural networks, but required extensive gradient-based training [26, 27, 11]. Another important difference is that our resulting neural network was composed solely of place cells, although neuromodulated, and no other types of neurons were present. This distinction is justified by the partially different task protocol and internal architecture, which constrained the tuning dynamics and also did not receive visual information as in [25]. Furthermore, our model relied on predefined grid cell layers, which constituted a strong and sufficient inductive bias, and did not have to be learned from scratch.

An additional relevant aspect is the consideration of the place cell layer as an explicit graph data structure, on which the path-planning was applied, meant to be a neural-based variant of the Dijkstra algorithm [90], a popular choice for graph computations. In fact, our algorithm relied on biologically plausible neural operations, such as activity propagation through the network and the utilization of synaptic activity traces as masks to identify the best neighboring place cells. This structured representation introduced a strong inductive bias, promoting robustness and flexibility across environments with varying layout complexity. Moreover, this lifted the need to learn an approximation of it through network dynamics and rely on a greater variability in neural receptive fields.

A similar approach using boundary cells has also been used for applying the successor representation (SR) framework to large environments [91], with the emergence of place and grid cell tuning. More broadly, on the one hand, several aspects of the SR framework align with our models: the distinction of the state map and the reward/boundary signals, and the use of an update rule inspired by temporal difference learning [31, 32, 33]. In addition, several studies have investigated biologically plausible implementations [34, 36, 35], obtaining good and promising matches with experimental results. On the other hand, the construction principles of our architecture required fewer assumptions about the state space size (the environment area can be almost arbitrary large), while learning the basis of the state representation (place cells) and the successor matrix (associated to the lateral connectivity) is more direct and immediate than in the standard SR formulation. These modeling choices reflect our interest in the one-shot generation of an explicit place cell map with an associated graph representation, while the SR may also produce non-spatial neural profiles. Finally, our architecture was better suited to integrate and study our formulation of neuromodulated plasticity, given direct access to a spatial map.

Adaptability was tested by occasionally moving the reward position, leading to the generation of an internal prediction error that was used to update its representation on the map. The agent was proved capable of unlearning previous associations, returning to exploration, and memorizing new reward locations. This behavioral protocol is similar to previous work [92], in which dopaminergic and cholinergic activity was utilized within a Hebbian plasticity rule to strengthen or weaken reward-associated spatial representations. However, alternatively to exploiting neuromodulators with opposite valence, we followed the direction of temporal-difference learning and predictive coding, a direction linked to hippocampal representations [29, 93] and explored various computational approaches [94, 95, 96]. This choice departed from our focus on using the data encoded in the cognitive map itself, in which each position has an associated neuromodulation value array, encapsulated in the modulator connection weights, that functions as expected sensory experience to compare with the actual sensory experience and obtain a learning signal. This mechanism aligns with long-standing theories linking neuromodulation, particularly dopamine, to prediction error signaling [97, 98, 48, 57, 60, 41].

However, a limitation of our current implementation is the simplification of neuromodulation to externally defined sensory events represented as a twodimensional input. Furthermore, neuromodulation in our model only influences value representation and place field modulation, which captures only a subset of the broader roles neuromodulators play in the brain.

Lastly, the relevance of active modulation of the neuronal properties of place cells was confirmed through simulated ablation experiments. The primary finding was the statistically significant role of active boundary modulation, obtained by comparing it with a model variant with disabled neuromodulation. This supports the importance of having a flexible representation of the environment layout.

Further investigation involved testing the effect of density and gain modulation separately. The results reported a significant impact of altering the density of place cells on the total count of collected rewards. In general, these results are consistent with the experimental observations of alteration of place cells following reward events [50, 99], in particular in terms of increased clustering of cells [100, 101], reminiscent of changes in firing rate after contextual changes [83, 84]. This relocation mechanism was stronger for the boundary cells, with the trend of pushing the place cell centers closer to the boundaries, and it proved to be statistically significant in improving performance. For reward cells, the effect was weaker and did not improve behavior.

Concerning the modulation of place fields, there is significant experimental evidence of their alternation during reward events [85, 102, 86], some reporting shrinkage near reward objects [81], and more markedly near boundaries [82]. In this direction, two possible experimental predictions can be made, based on the constraint of using self-motion cues for the generation of cognitive maps. Firstly, the size of the place fields at the boundaries is initially homogeneous, and only after meaningful events is it reshaped, such as after collisions. This can be tested by correlating place cells with the count of relevant events occurring near you. Secondly, it can be done similarly for rewarding events, and possibly going further by blocking dopamine release in order to assess its causal involvement.

In our setup, the place-field modulation was implemented by scaling the neural activation gain. This led to a general reduction in field size across all modulated cells. In the case of boundary cells, this shrinking effect may account for the improved performance in two ways: first, by increasing the spatial resolution of the environment layout [103]; and second, by freeing up space for other place cells to form near. The latter may result from reduced lateral inhibition and the inward shift of the field center toward the boundaries. However, gain modulation alone did not produce statistically significant effects, indicating that it must be coupled with density modulation to produce measurable improvements.

For reward place cells, the impact of gain modulation was less pronounced. This may be due to the simplicity of the experimental protocol or reward signal, with the latter being defined purely in spatial terms and lacked any meaningful non-spatial feature.

Previous studies have proposed that Locus Coeruleus (LC) is involved in place cell reorganization [49] and overrepresentation near reward locations through the co-release of dopamine and noradrenaline [51]. A possible experimental prediction that emerges from our results is that preventing place cell density modulation may indirectly decrease the quality of reward-directed navigation. More specifically, this can be tested in animal models by pharmacological blocking of LC afferent connections to CA1, once reward has been experienced. Then, the performance can be evaluated by considering: the count of rewards fetched; the difference between the animal’s trajectory between the initial and reward location and the theoretical spatial geodesic (shortest path); and the average distance from the walls. However, a relevant confounding variable in this set-up is the difficulty of isolate CA1 inputs, and the involvement of multiple and complex additional signals in the generation of the cognitive map.

In conclusion, this work showed a possible architecture for coupling emergent spatial representations with neuromodulated plasticity to achieve an experience-driven cognitive map. The reliance on some spatial and algorithmic inductive biases, grid cells, and a planning algorithm supports the idea of a label graph for goal navigation. Future work can investigate the application to other spatial domains, such as motor control and three-dimensional navigation. In addition, a richer input feature can be added, such as visual information [104], as well as new neuromodulators that encode different sensory dimensions or internally generated signals.

## Acknowledgements & Statements

The authors declare no competing interests.

The code is publicly available and can be found at https://github.com/iKiru-hub/PCNN.

This research was funded by the European Union’s Horizon 2020 research and innovation programme under the Marie Skłodowska-Curie grant agreement Nº 945371 and the University of Oslo.

The research presented in this paper has benefited from the Experimental Infrastructure for Exploration of Exascale Computing (eX3), which is financially supported by the Research Council of Norway under contract 270053.

## 5. Appendix

### 5.1 Grid cell module

Describe a correspondence between the global environment in which the agent moves, a two-dimensional Euclidean space **R**^2^, and a bounded local space of a grid module, corresponding to a torus.

The global velocity **v** = {x, y} is then mapped to a local velocity, scaled by a speed scalar 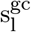 specific to the grid cell module l, which determines its periodicity in space.

The choice of a toroidal space is motivated by consolidated experimental evidence of the neural space of grid cells, which are organized in modules of different sizes spanning the animal’s environment. However, the shape of their firing pattern is known to be hexagonal, which corresponds to the optimal tiling of a two-dimensional plane, giving rise to a neural space lying on a twisted torus. In this work, for simplicity, we consider a square tiling and thus a square torus, without much loss of generality except for the slight increase of grid cells required for a sufficiently cover.

A grid cell module l of size N^gc^ is identified by a set of positions defined over a square centered on the origin and size of 2, such that {(x_i_, y_i_) | i ∈ N^gc^ ∧x_i_, y_i_ ∈(–1, 1)}. This local square space has boundary conditions for each dimension, such that, for instance, when 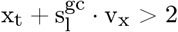 the position update is taken to the other side 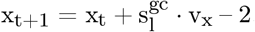, where 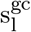 is the scale of the velocity in the local space of the module l with respect to the real global agent velocity v = {v_x_, v_y_}. When the module is initialized, the starting positions of its cells are uniformly distributed over the square forming a lattice. When the agent is reset in a new position at the beginning of new trial, a displacement vector is applied to the last cells positions such that the mapping between the module local space and the global environment is preserved.

The firing rate vector of each cell is determined with respect to a 2D Gaussian tuning curve centered at the origin at (0, 0), and it is calculated as 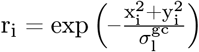, where 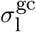 is the width of the tuning curve for module l. An illustration of the receptive field in a 2D environment and a toroidal space is reported in Figure 5**a**-**b**.

**Figure 5.**
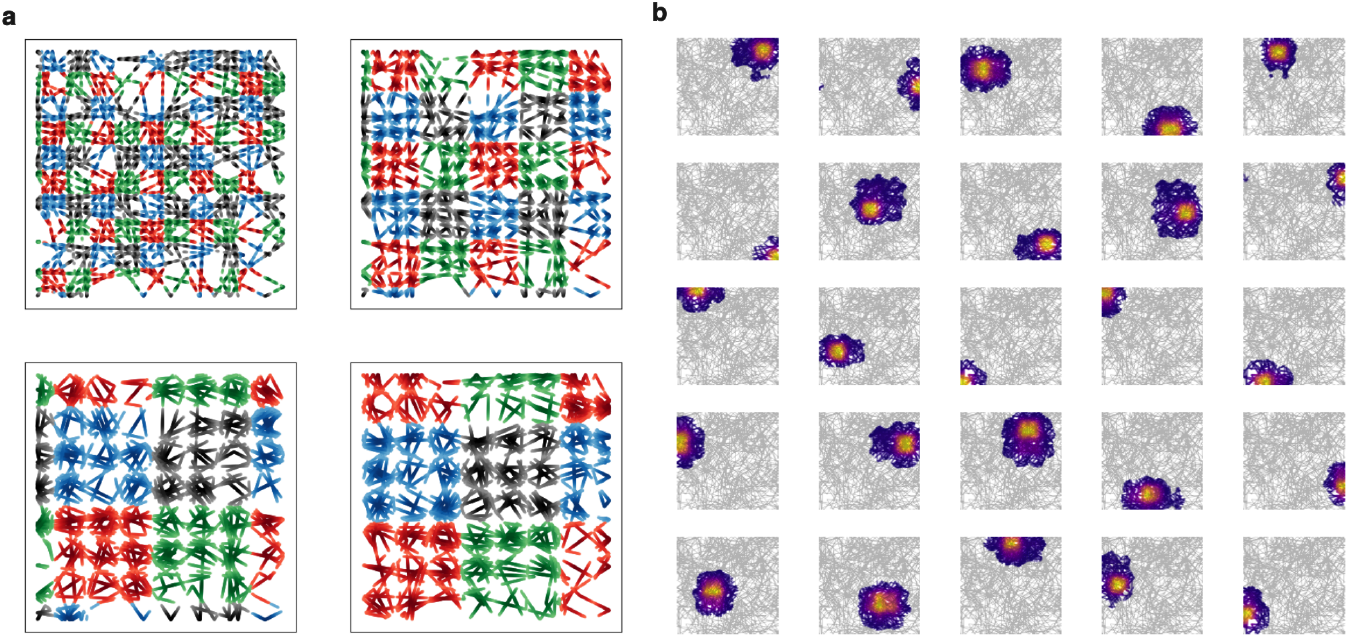
Place fields obtained from grid cells activity. **a**: *grid cell modules with different granularity represented over a continuous trajectory in an open space. For visualization purposes, each module is represented as composed of four sub-modules of 9 grid cells each, whose periodic tuning generates activity that repeats in space*. **b**: *place cells whose spatial tuning has been obtained from the concatenation of the grid cells population vector*.

The final population vector of the grid cell network GC is the concatenated and flattened firing rate vector of all modules **u**^GC^.

In our model, each grid cell had a tuning width of 0.04. They were defined as 8 modules of size 36, and the relative speed scales were

{1., 0.8, 0.7, 0.5, 0.4, 0.3, 0.2, 0.1, 0.07}.

#### 5.2 Place cells

##### Tuning formation

The tuning of a new place cell is simply defined as the current GC population vector 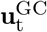, and its index is that of the first silent cell, which is added to the forward weight matrix 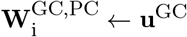.

In order to avoid overlapping of place fields, lateral inhibition is implemented. More specifically, the tuning process is aborted in case the cosine similarity of the new pattern and the old ones is greater than a threshold 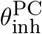.

Each cell represents a position in the GC activity space, which can be considered a node within a graph of place cells (PC). Although it is totally possible to only use the N^GC^-dimensional tuning patterns and be agnostic about the dimensionality of the space in which the agent lives, to simplify the calculations, we mapped each pattern to 2D positions in a vector space. Then, the PC recurrent connectivity matrix is calculated with a nearest neighbors algorithm, which instead of a fixed number K of neighbors uses a lateral distance threshold 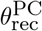.

##### Activity

The current firing rate of the PC population is determined by the cosine similarity between the GC input and the forward weight matrix, then passed through a generalized sigmoid *ϕ*(z) = [1 + exp(–*β*(z – *α*))]^–1^. The parameter *α* represents the activation threshold, or horizontal offset, while *β* the gain, or steepness.

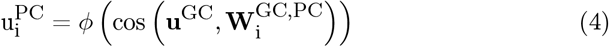

It is also defined as an activity trace, which has an upper value of 1 and decays exponentially:

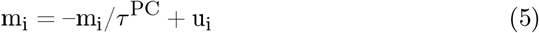

It is used as a proxy for a memory trace.

In the model, a PC population is defined by its average place field size, determining the granularity of the representation of the place. In plot 5**b** it is illustrated an example of place cells layer tuning obtained from a continuous trajectory over a square environment.

#### 5.3 Modulation

Neuromodulation is implemented as a reward-sensitive signal, represented as DA (imitating the function of dopamine), and a collision-sensitive signal, represented as BND (for boundary). Its dynamics are defined in terms of a leaky variable v whose state is perturbed by an external input x, whose qualitative meaning differs for each neuromodulator k.

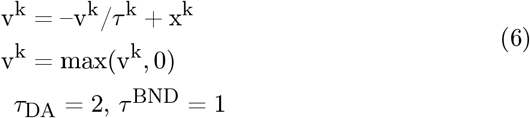

##### Learning rule

The connection weights **W**^k^ are updated according to a plasticity rule composed of a Hebbian term, involving the leaky variable, the place cells that are above a certain threshold *θ*^k^, and the current connection weights value:

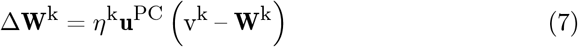

where the weight contribution of the Hebbian update and *η*^k^ is the learning rate: *η*^DA^ = 0.9, *η*^BND^ = 0.9. Additionally, connections values are kept nonnegative.

##### Active neuronal modulation

Neuromodulation acts on the neuronal profile of the place cells by affecting the value of the activation gain and relocates the center of their tuning.

Gain modulation is implemented using activity traces and a constant reference gain value 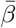:

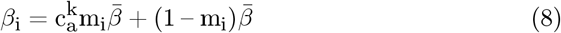

where 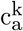 is a scaling gain parameter, and if it is 1 then no modulation occurs. Concerning center relocation, it is applied to recently active neurons with non-zero trace m_i_. For a place cell i with position 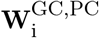 (in the grid cell space), a displacement vector q_i_ is calculated with respect to the current position **u**^GC^:

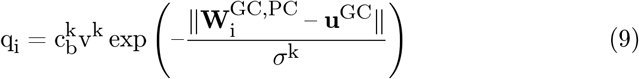

where 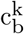 is a scaling relocation parameter, while *σ*^k^ is the width of the Gaussian distance. This displacement is used to move in the GC activity space and get the new GC population vector to use as a tuning pattern.

For this modulatory action, they were considered only cells whose activity trace was greater than the modulator-specific threshold *θ*^k^, to prevent the involvement of silent cells. In this case, it is also ensured that the center of the new place field is at a minimum distance 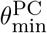 from the others; here, Euclidean distance is used.

#### 5.2 Decision making

##### Behaviour selection logic

The possible behaviors are *exploration* and *exploitation*, and an action is defined as a 2D velocity vector. For exploration, an action can be generated either as random navigation, using a polar vector of fixed magnitude (the speed) and angle sampled from a uniform distribution, or as a step within a goal-directed navigation plan to reach a random destination, which corresponds to a randomly sampled existing place cells. During goal-directed navigation, the magnitude of the velocity vector is less than or equal to a fixed speed value, depending on the distance from the next target position in the plan. Instead, for the exploitation behavior, the action is a step within a goal-directed navigation towards the reward location. The behavior selection process depends on the experience of collisions, the presence of a plan, and the success of navigation planning. A diagram of this logic is shown in Figure 6.

**Figure 6.**
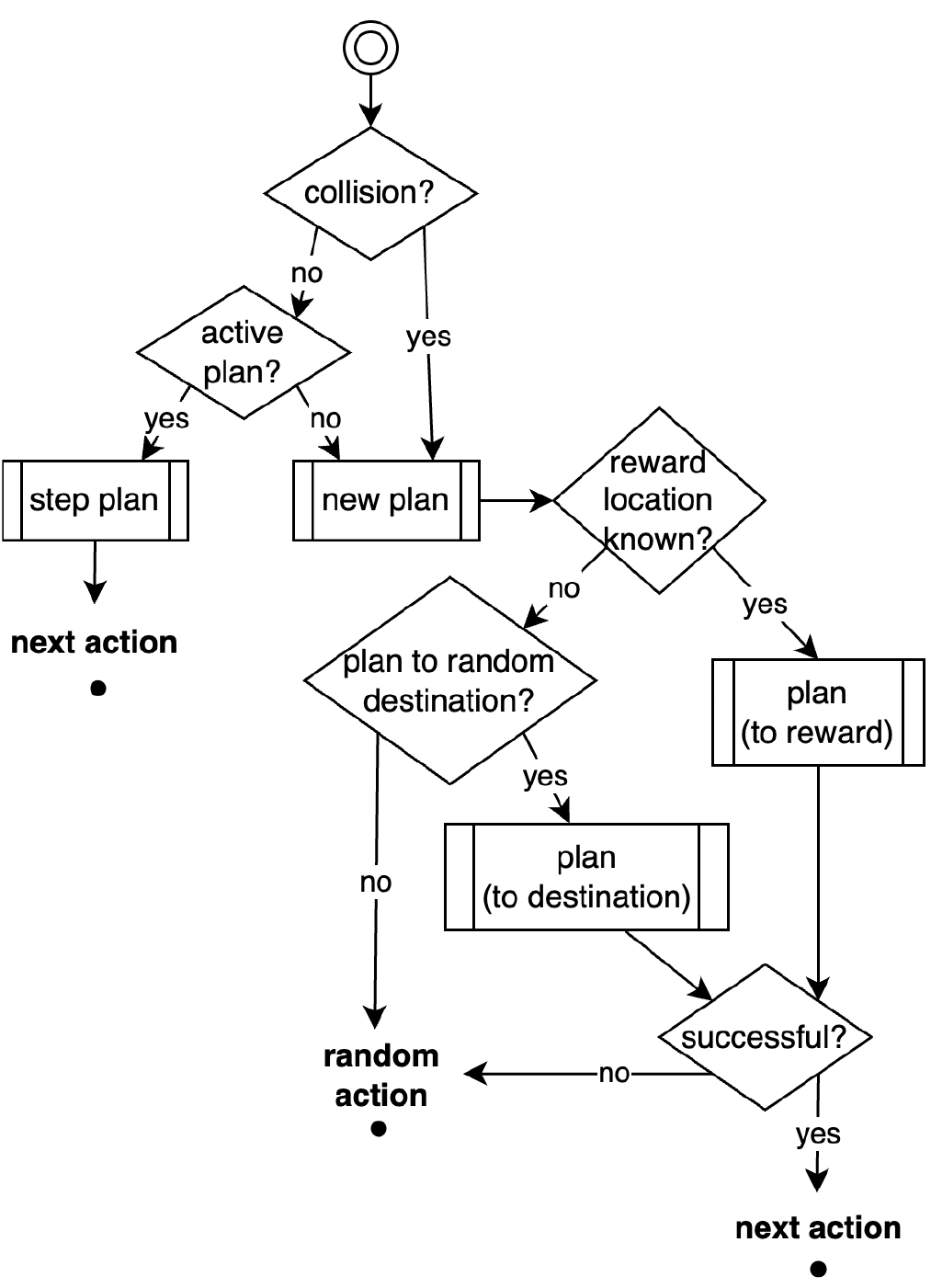
Diagram of the behaviour selection process.

The positions of the agent and of the target location for planning are identified by the place cells population vector. In particular, the reward position (x_r_, y_r_ is determined by the weighted average of the centers x_i_, y_i_ of the place cells with respect to their DA-modulated connections weights. Furthermore, only the top five place cells are considered.

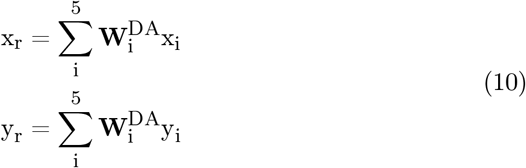

##### Path-planning algorithm

The planning of a new route is implemented as a path-finding algorithm based on the place cell graph, provided as a connectivity matrix C. Its particularity is the use of a weighting 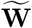 of the nodes according to the neuromodulation map. A description is reported in algorithm 1.

###### Algorithm 1

Activity-based Pathfinding

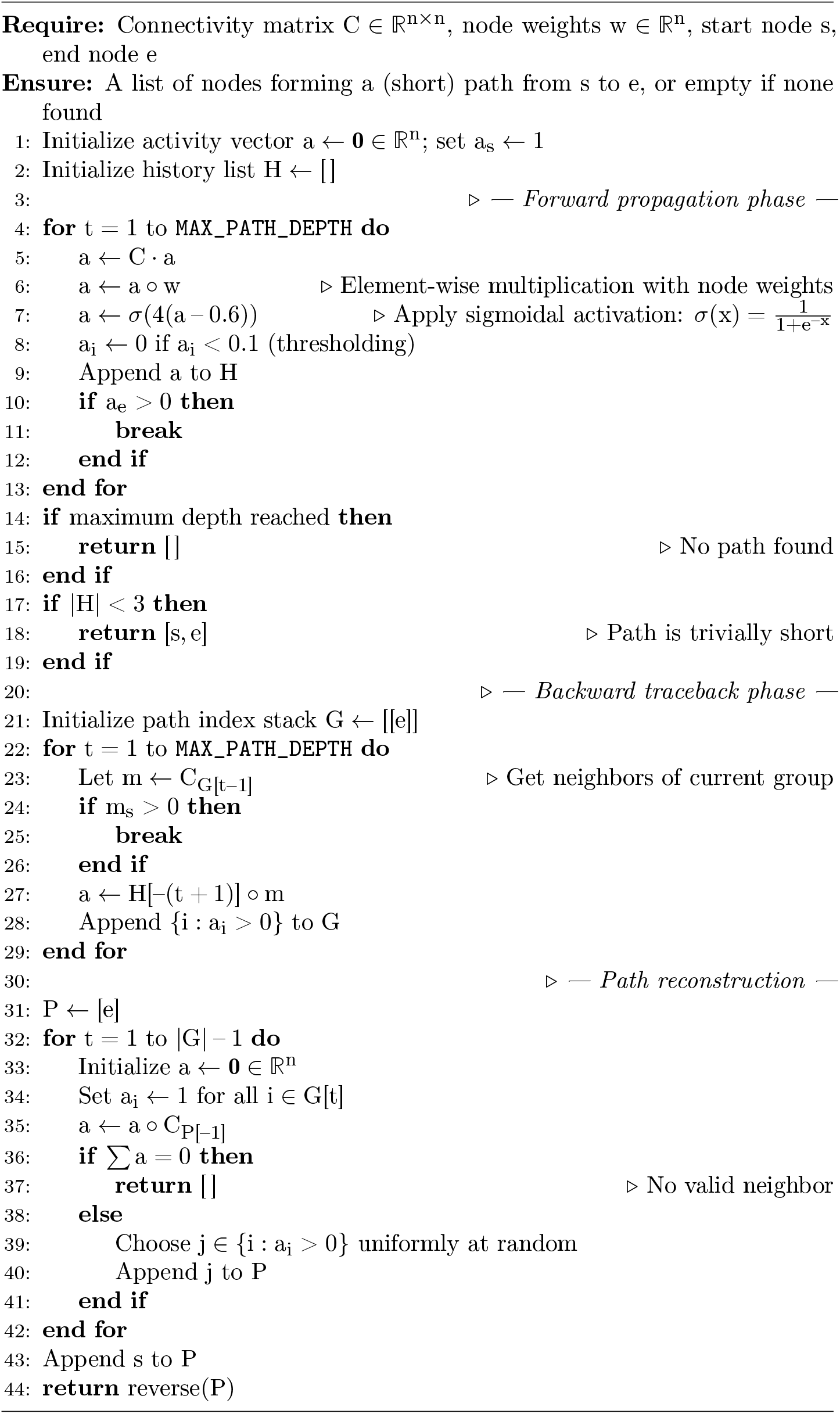

##### Comparison with previous architectures

Several previous computational models show structural and conceptual similarities with the present work. A prominent category among them employs deep neural networks, often with recurrent components, and is based on gradient-based learning strategies such as backpropagation over time. These models typically require multiple training episodes or large datasets for convergence. In contrast, our model adopts biologically inspired local plasticity rules and requires only a single training episode for adaptation.

Other models utilize spiking neurons [92] or explicit neural representations [29], and incorporate online learning rules more closely aligned with ours. These models also focus more directly on goal-directed navigation, in contrast to purely path-integration tasks. However, both of these rely on external spatial coordinates to represent the current position. Our model instead constructs an internal coordinate system by integrating its own velocity output, enabling endogenous spatial tracking.

A summary of such comparison is reported in table 1.

**Table 1:**
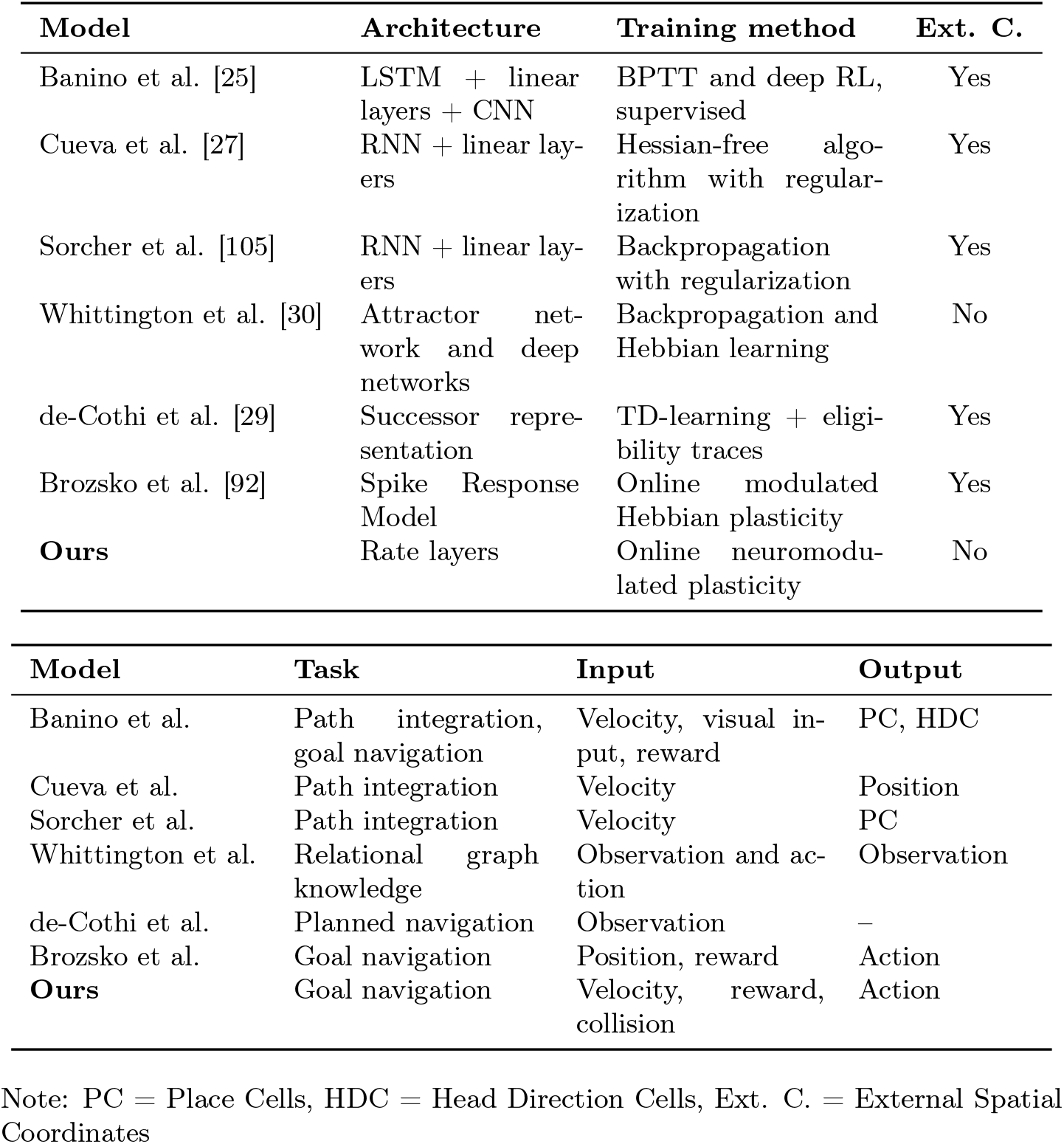
Comparison of neural network models for spatial navigation and representation

| Model | Architecture | Training method | Ext. C. |
| --- | --- | --- | --- |
| Banino et al. [25] | LSTM + linear layers + CNN | BPTT and deep RL, supervised | Yes |
| Cueva et al. [27] | RNN + linear layers | Hessian-free algorithm with regularization | Yes |
| Sorcher et al. [105] | RNN + linear layers | Backpropagation with regularization | Yes |
| Whittington et al. [30] | Attractor network and deep networks | Backpropagation and Hebbian learning | No |
| de-Cothi et al. [29] | Successor representation | TD-learning + eligibility traces | Yes |
| Brozsko et al. [92] | Spike Response Model | Online modulated Hebbian plasticity | Yes |
| **Ours** | Rate layers | Online neuromodulated plasticity | No |

| Model | Task | Input | Output |
| --- | --- | --- | --- |
| Banino et al. | Path integration, goal navigation | Velocity, visual input, reward | PC, HDC |
| Cueva et al. | Path integration | Velocity | Position |
| Sorcher et al. | Path integration | Velocity | PC |
| Whittington et al. | Relational graph knowledge | Observation and action | Observation |
| de-Cothi et al. | Planned navigation | Observation | – |
| Brozsko et al. | Goal navigation | Position, reward | Action |
| **Ours** | Goal navigation | Velocity, reward, collision | Action |

Note: PC = Place Cells, HDC = Head Direction Cells, Ext. C. = External Spatial Coordinates

#### 5.5 Environments

The game in which the model is tested has been developed with the Pygame Python library, used under the GNU LGPL version 2.1 license and available at https://github.com/pygame/pygame. The layout of the environment consisting of a customizable arrangement of vertical and horizontal hard walls with variable length and fixed width. In Figure 7, some samples are shown below.

**Figure 7.**
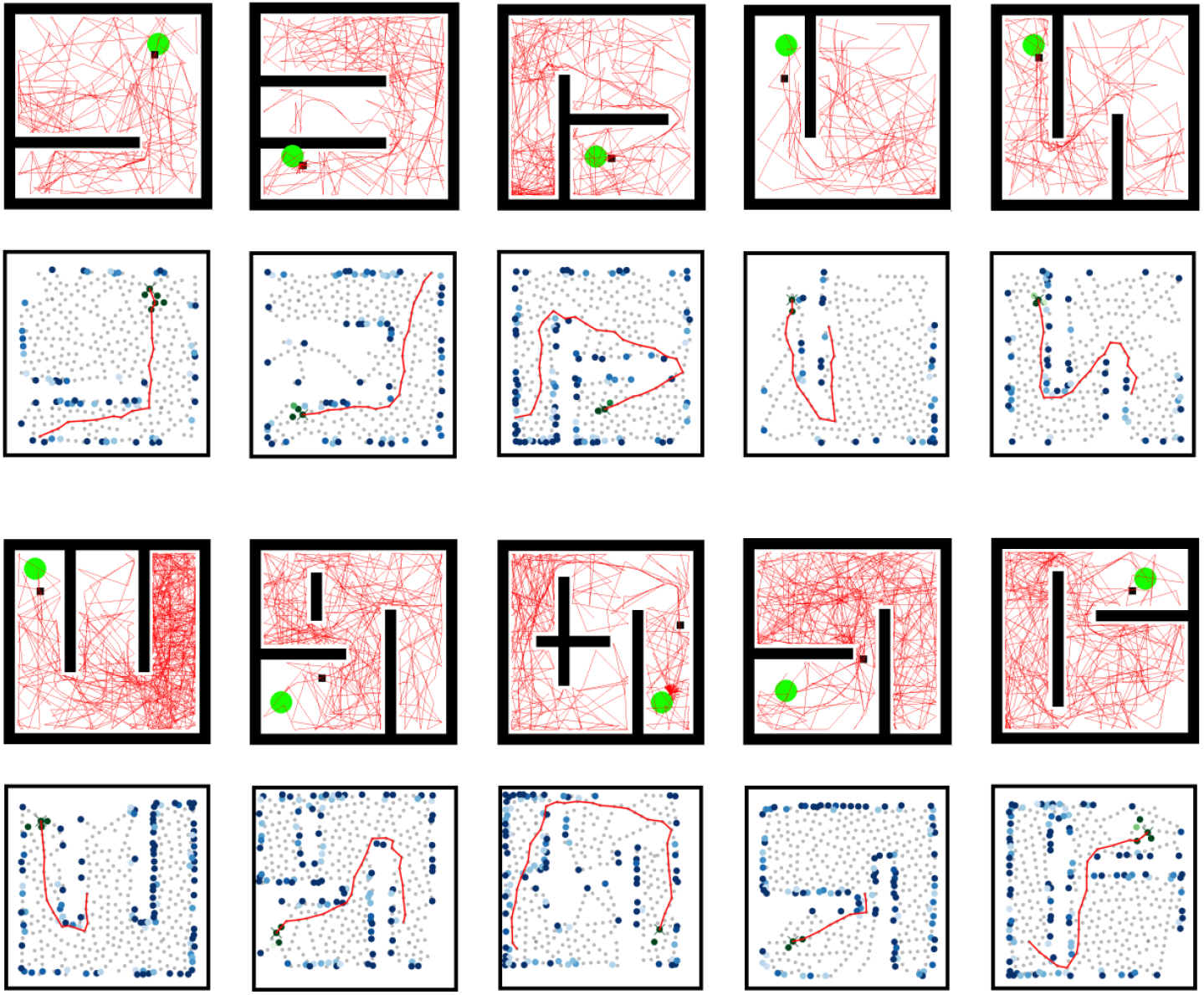
Sample of generated environments.

The reward object is defined as a circle with a size the 5% of the total environment area. When the agent’s position is within its boundary, a binary signal R*∼* ℬ(p_r_) is provided drawn from a Bernoulli with probability p_r_ = 0.6. The duration of the reward fetching is set to 2 time steps.

The agent object is defined as a square with a size of 3. 3% of the total area of the environment.

The testing protocol was inspired by the behavior of animals that venture into new territories in search of food. It was divided into two parts:

- **exploration phase**: the agent was placed in a random location within the environment for 10,000 time steps. In this phase, the reward is not present. Further, in order to force greater exploration of the environment, every 3000 steps it was teleported to another random location. This external intervention was intended to mitigate randomness in the agent’s exploratory behavioral strategy.
- **reward phase**: a reward is inserted in a random location and available to be discovered. When it is encountered, the agent is teleported to a random location within the environment, and after a fixed about 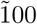 time step it is enabled to use reward seeking behavior, in the form of goal-directed navigation. The total duration of this phase is set to 20’000 time steps.

An episode is defined as a continuous trajectory during the reward phase, namely a set of time steps starting from when the agent is placed in a position until either it finds the reward or the simulation ends.

##### Detour experiment

The protocol is modified such that after a fixed number of episodes the layout of the environment is changed, *e*.*g*. a wall is inserted. This experiment is meant to test the ability to reach the reward location by using the same cognitive map, and possibly update it with the new sensory information, such as the detection of the new boundaries.

##### Changing reward experiment

During the reward phase, the reward location is changed after a fixed number of fetches.

##### Optimization

The model hyper-parameters such as the constants for the neural dynamics and the behavior selection have been optimized through an evolutionary algorithm. Initially, an initial population of individuals with different random genomes (string of hyperparameter values) is sampled and evaluated. Then, the population of a new generation is constructed from the first by combining and mutating the genomes of the top-ranked individuals from the previous generation. More in detail, for the sampling of the new generation, we used the Covariance Matrix Adaptation algorithm, in which the shape of distribution of genome values is iteratively adapted according to the recent performances. The fitness used to evaluate the individuals was a tuple consisting of the number of rewards collected and the number of collisions. In particular, the latter was calculated starting from the time when the reward position was first discovered; in this way, at least the stochasticity of the exploratory behavior was excluded.

The hyperparameters evolved are: neural gain *β*, lateral inhibition threshold 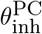, lateral distance threshold 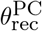, activity trace time constant *τ* ^PC^, reward modulation scale 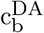, reward modulation spread 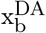, boundary modulation scale 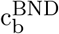, boundary modulation spread 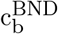, reward gain modulation 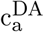 and boundary gain modulation 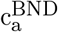.

